# Autism spectrum disorder risk genes have convergent effects on transcription and neuronal firing patterns in primary neurons

**DOI:** 10.1101/2025.03.25.645337

**Authors:** Alekh Paranjapye, Rili Ahmad, Steven Su, Abraham J. Waldman, Jennifer E. Phillips-Cremins, Shuo Zhang, Erica Korb

## Abstract

Autism spectrum disorder (ASD) is a highly heterogenous neurodevelopmental disorder with numerous genetic risk factors. Notably, a disproportionate number of risk genes encode transcription regulators including transcription factors and proteins that regulate chromatin. Here, we teste the function of nine such ASD-linked transcription regulators by depleting them in primary cultured neurons. We then define the resulting gene expression disruptions using RNA-sequencing and test effects on neuronal firing using multielectrode array recordings. We identify shared gene expression signatures across many ASD risk genes that converge on disruption of critical synaptic genes. Fitting with this, we detect robust disruptions to neuronal firing throughout neuronal maturation. Together, these findings provide evidence that loss of multiple ASD-linked transcriptional regulators disrupts transcription of synaptic genes and has convergent effects on neuronal firing that may contribute to enhanced ASD risk.

## Introduction

Autism Spectrum Disorder (ASD) is a highly prevalent neurodevelopmental disorder with numerous risk genes (De Rubeis et al. 2014; Iossifov et al. 2014; Sullivan and Geschwind 2019; de la Torre-Ubieta et al. 2016). Major functional groups of ASD risk genes have emerged including a large portion that encode proteins that regulate transcription (Iossifov et al. 2014; O’Roak et al. 2012; Parikshak et al. 2013; De Rubeis et al. 2014). Of these ASD-linked transcriptional regulators, many encode proteins that regulate chromatin, the complex of DNA and histone proteins that help to regulate transcription. Others encode transcription factors while still others directly modify DNA itself. Thus, distinct groups of proteins with disparate functions in transcriptional regulation can lead to overlapping phenotypic outcomes.

We previously tested the effects of disrupting a set of ASD risk genes that function as chromatin regulators and found that they affected expression of a common set of genes that encode synaptic proteins (Thudium et al. 2022). This gene expression signature was detected in multiple systems and, together with the broader literature (Satterstrom et al. 2020; Zhao et al. 2018), suggest that neurons have gene sets that are highly susceptible to chromatin disruptions regardless of the specific manipulation with similar signatures found across other psychiatric disorders (Rajarajan et al. 2018; Schrode et al. 2019). Further, numerous studies have identified sets of synaptic genes that have unique chromatin features and that are disrupted in ASD, either directly as ASD risk genes or indirectly following disruption of ASD-linked transcriptional regulators (Satterstrom et al. 2020; Zhao et al. 2018). Both our work and others (Thudium et al. 2022; Satterstrom et al. 2020) found that the targets of ASD-linked transcriptional regulators do not directly regulate other ASD-linked gene more than is expected by chance and instead target synaptic genes more broadly. However, prior analysis was limited to a handful of chromatin regulators. Whether similar transcriptional signatures are detected in response to disruption of transcriptional regulators more broadly beyond just chromatin-modifying enzymes, and whether such changes result in shared functional outcomes remain unclear.

Here, we sought to address these outstanding questions, using a primary neuronal culture model to allow for comparisons within a highly controlled, genetically identical population of neurons. We then defined shared transcriptional and functional outcomes of distinct ASD-linked genes during neuronal maturation to improve our understanding of the molecular basis for ASD and related neurodevelopmental disorders.

## Results

### Depletion of ASD-linked transcriptional regulators affects gene expression

Here, we sought to define shared gene expression signatures amongst multiple ASD-linked transcriptional regulators including ASH1L, CHD8, DNMT3A, KDM6B, KMT2C, MBD5, MED13L, SETD5, and TBR1. We selected these targets to jointly test the roles of chromatin regulators, DNA modifying enzymes, and transcription factors, expanding upon prior work that focused on chromatin regulators alone. Importantly, here we selected chromatin modifying enzymes that target different both distinct (KDM6B, KMT2C) and overlapping (ASH1L, SETD5) histone sites, histone remodelers from 2 different complexes (CHD8, MED13L), a DNA modifying enzyme (DNMT3A), a transcription factor (TBR1), and a non-catalytic chromatin-complex protein (MBD5) (**Table 1**). Additional criteria included that all are high confidence ASD genes (scored as a ‘1’ within the SFARI gene module and as a significant TADA gene), all lead to well-defined syndromes or associated phenotypes caused by loss-of-function mutations or deletions, and all have been successfully modeled in mice (Fu et al. 2022; Gao et al. 2021; Shen et al. 2019; Brauer et al. 2023; Lavery et al. 2020; Nakamura et al. 2024; Moore et al. 2019; Beighley et al. 2020; Bernier et al. 2014; Hurley et al. 2021; Adegbola et al. 2015; Christian et al. 2020; Li et al.; Huang et al. 2019; Mullegama and Elsea 2016; Camarena et al. 2014; Chatterjee et al. 2023).

**Table 1:** Function and association data for the nine transcriptional regulators chosen for analysis. SFARI gene score of 1 indicates high confidence for implication in autism spectrum disorder. Evaluation of Autism Gene Link Evidence (EAGLE) score ranges from 6 (limited) to 12+ (definitive) roles in autism for validated targets. Dashes indicate no known disorder or calculated EAGLE score.

To test these genes of interest, we used primary neuronal cultures derived from E16.5 embryonic mouse cortical tissue to generate a highly pure neuron population (**Fig. 1A**) (Thudium et al. 2022). This allows for all targets to be tested within the same cell population without the complexity and heterogeneity of brain tissue. Further, this system provides a genetically identical background which avoids confounding variables from the mixed backgrounds from patient samples. This system is also sufficiently tractable to test multiple candidate genes in parallel within neurons derived from the same embryo, thus providing a highly controlled system for comparisons and true biological replicates. Lastly, this system allows for targeting of transcriptional regulators in neurons following isolation from tissue and establishment of neuronal identity. This ensures that the same cell types are measured across conditions by avoiding disruptions to neuronal precursor cells that ultimately might affect cell identity, or the ratio of inhibitory and excitatory neurons present during testing. While disruptions during early neurogenesis are almost certainly highly relevant to the biology underpinning ASD, here we specifically sought to focus on effects within post-mitotic neurons to allow for direct comparisons in equivalent cell types without disrupted developmental trajectories. Together, this system therefore allows for parallel testing of multiple genes across multiple biological replicates on a genetically identical background during a key period of neuronal maturation without changes in cell identity.

**Figure 1:**
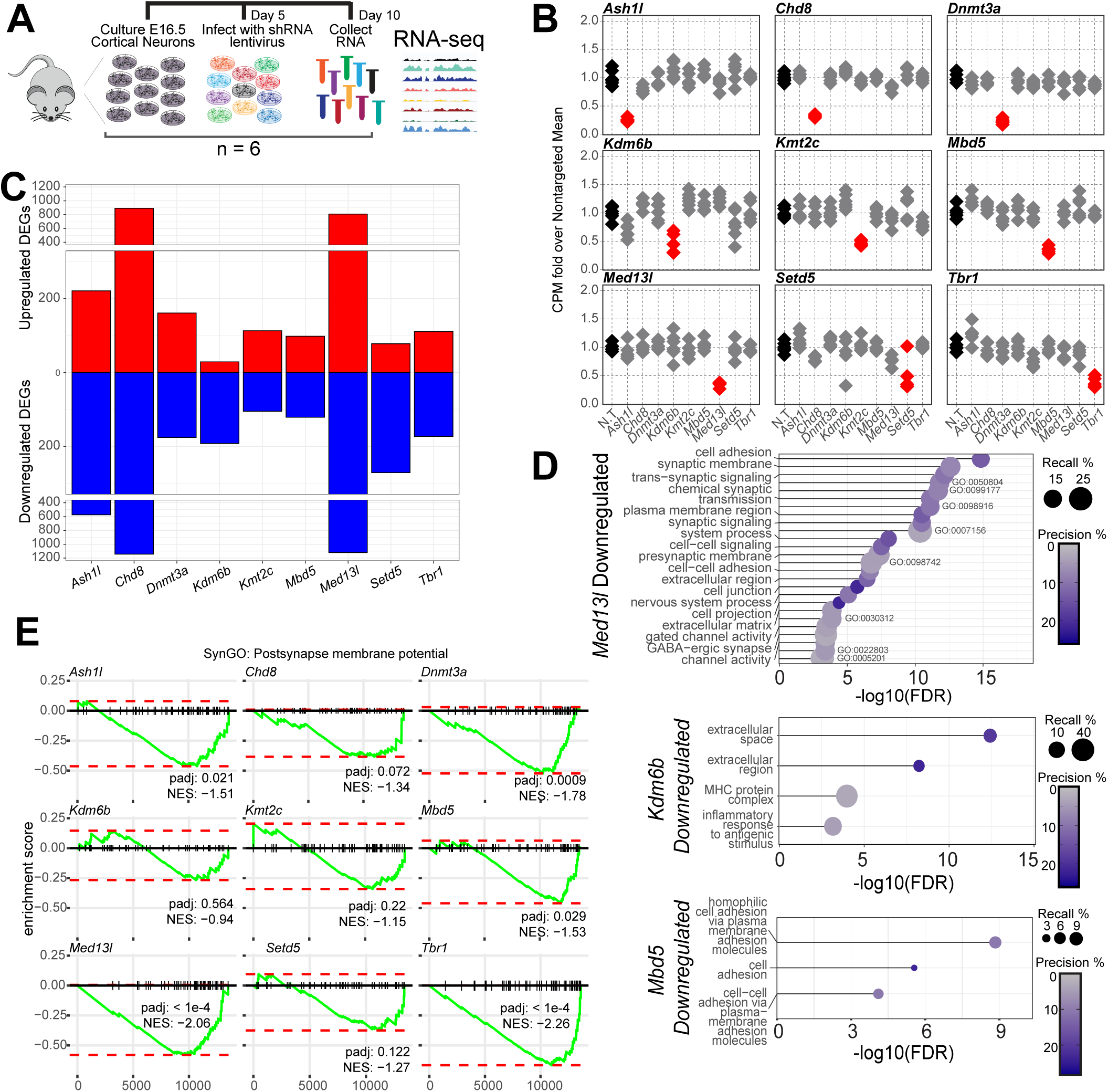
Gene expression analysis of nine independent chromatin modifier depletions in primary mouse neurons. A. Schematic of the experimental timeline for the comparison of transcriptomes between chromatin modifier depletions. B. Counts per million (CPM) for the nine chromatin modifiers following lentivirus-mediated shRNA depletion of each target, relative to the average of nontargeting (N.T.) treated neurons (n = 6). Red shows significant changes. Gray indicates that no others pass statistical significance by the DESeq2 negative binomial distribution with multiple testing correction. C. Total up or downregulated differentially expressed genes (DEGs) in the pairwise comparison of each depletion versus N.T. treated neurons. Full DEG lists shown in Supplemental Table 3. D. Gene ontology analysis of significantly downregulated gene sets. Recall is the proportion of functionally annotated genes in the query over the number of genes in the GO term. Precision is the number of genes found in the GO term over the total number of genes in the query. Results for other targets shown in Supplemental Table 4. E. GSEA of genes in the SynGO postsynaptic membrane potential term for each depletion transcriptional signature. The green line is the running, normalized enrichment score across all genes ranked by their change in expression from most increased at 1 to least increased. The red dotted lines specify the maximum and minimum running enrichment score per depletion. NES indicates the directional normalized enrichment score across all genes.

We cultured neurons derived from 6 wild-type E16.5 cortices (3 male and 3 female) and infected neurons with lentivirus containing shRNA at 5 days in vitro (DIV) to achieve partial depletion of targets and model a partial loss of function variant. Depletion of each target transcript was confirmed at DIV 10 (**Fig. S1A**) and, where antibodies were available, similarly confirmed depletion at the protein level here (**Fig. S1B**) or in prior work (Thudium et al. 2022). We then performed RNA-sequencing at DIV 10, the timepoint when we previously identified robust transcriptional effects of multiple ASD-risk genes (Thudium et al. 2022). We again confirmed knockdown of each target, and, similar to prior findings (Thudium et al. 2022), determined that these transcriptional regulators do not directly affect gene expression of the other ASD-linked transcriptional regulators examined here (**Fig. 1B**). This suggests that any shared downstream effects are not simply due to one target causing disruption of another and thus also disrupting their target genes indirectly.

Next, we defined differentially expressed genes (DEGs) following depletion for all targets (**Fig. 1C, Supplemental Data Table 1**). Importantly, we detected minimal DEGs comparing non-targeting shRNA viral infection controls to non-infected neurons indicating that the viral infection alone did not perturb gene expression in neurons (**Fig. S1C-D**). All targeted transcriptional regulators caused gene expression changes as expected, although to markedly differing degrees. Notably, the degree of knockdown was not strongly correlated with the number of DEGs identified or the expression level of the target itself in this system (**Fig. S1E-F**), nor were the effects correlated with modifier expression in neurons isolated from mouse brains at similar times during maturation (**Fig. S1G-H**). This suggests that some ASD-linked transcriptional regulators have more robust effects on neurons during early maturation stages than others due their functional relevance to transcription in this culture system rather than due to technical factors. We also compared DEGs detected here and in separate experiments in prior work for the two overlapping targets (*Ash1l* and *Chd8*). Although datasets were collected over 7 years apart in time in different laboratories, and processed via different protocols and sequencing parameters, we did detect significant overlaps in gene expression changes (**Fig. S1I-J**).

To identify common functions encoded by disrupted gene, we performed gene ontology analysis on each group of significantly altered genes and found a wide range of enriched molecular and biological pathways across DEGs by depletion (Supplemental Data Table 2).

While not every gene set had significant gene ontology outputs, those that did predominantly included downregulation of genes associated with synapse formation and function (**Fig. 1D**) and upregulation of metabolism genes (**Fig. S2A**). For depletion conditions without these significant gene ontology terms, individual critical synaptic genes were still detected among downregulated DEGs (**Fig. S2B**). To determine the extent to which each depletion affected synaptic functioning beyond the most significant DEGs, we performed gene set enrichment analysis for each dataset examining expression change trends for postsynaptic membrane potential and synapse adhesion gene sets (**Fig. 1E, S2C**). 7/9 and 4/9 respectively showed significant downward trends for these key neuronal pathways. These analyses extend prior findings (Thudium et al. 2022) to other forms of transcriptional regulation beyond histone modifying enzymes. Interestingly, we found little overlap with high-confidence ASD and neurodevelopmental disorder (NDD) risk genes (**Fig. S2D**). This supports prior findings (Thudium et al. 2022) and suggests that ASD-linked transcriptional regulators cause neuronal disruption through mechanisms distinct from simply directly dysregulating other ASD-linked synaptic genes. Together, this work demonstrates that a broad range of ASD-linked transcriptional regulators serve to control expression of synaptic genes in neurons.

### Depletion of ASD-linked transcriptional regulators cause shared gene expression changes

Notably, each individual shRNAs can have off-target effects that may contribute to transcriptional changes. Therefore, we sought to define shared transcriptomic outcomes across targets to detect gene sets that are most sensitive to loss of ASD-linked genes rather than off-target effects of a single shRNA. To define overlapping transcriptional changes, we first directly compared DEGs from each target and found significant overlap in 63 of 81 comparisons (**Fig. 2A, Supplemental Data Table 3**). Further, we identified multiple genes shared by three or more conditions (**Fig. 2B-D)** and confirmed multiple shared genes by qRT-PCR (**Fig. S2E**). This fits with prior results that suggest that ASD-linked transcriptional regulators share multiple gene targets despite acting through disparate mechanisms in chromatin (Thudium et al. 2022).

**Figure 2:**
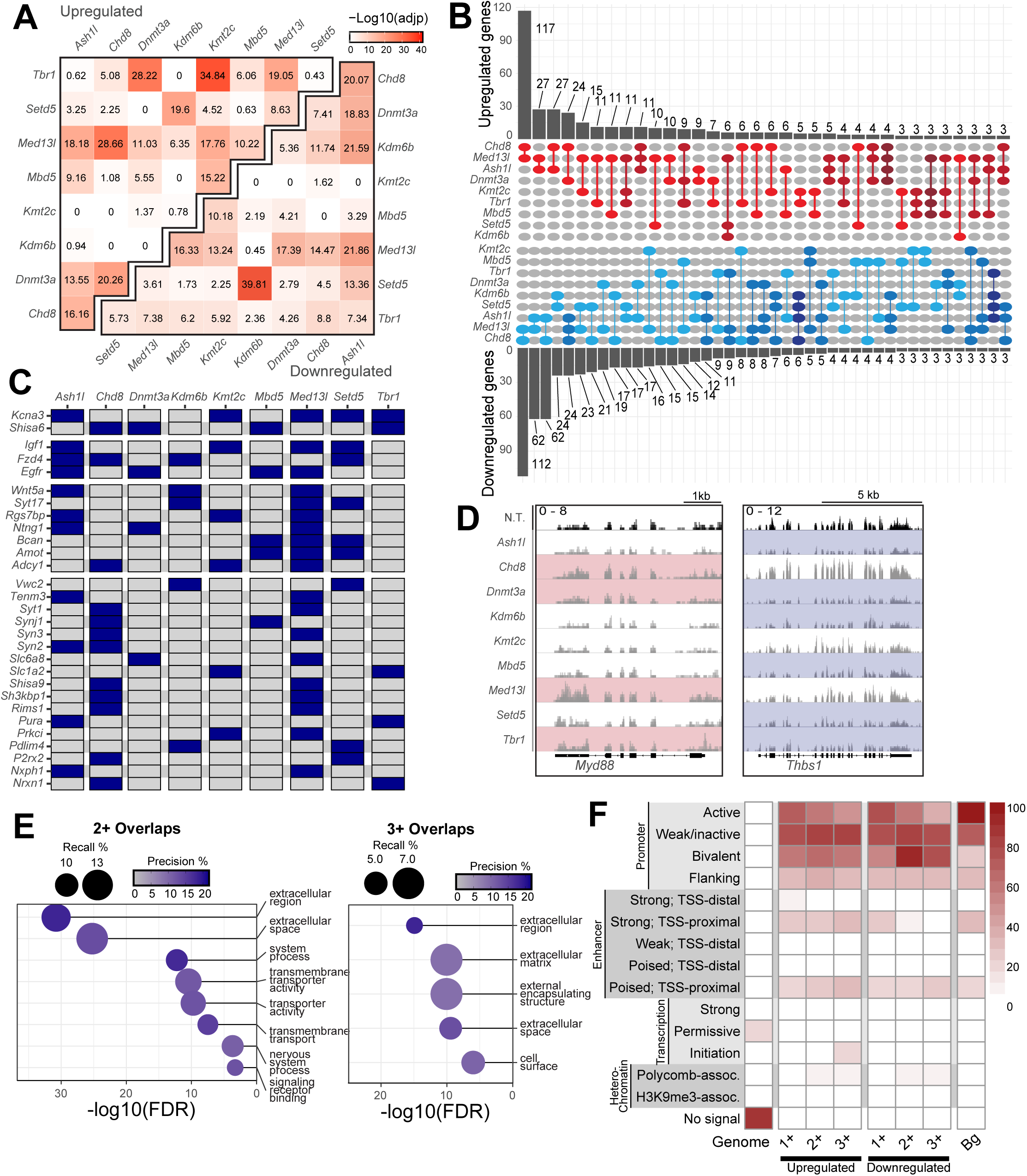
Identifying transcripts that co-dysregulated across depletions. A. Significance of overlap for up or downregulated DEGs after each depletion using hypergeometric tests. B. Upset plot of up or downregulated DEG found in at least two depletions. Gene placement prioritizes the largest intersection possible, only showing overlaps with 3 or more DEGs. The darker colors correspond to a greater overlap degree. C. DEGs that appear in at least two comparisons between depletion and N.T. control (in blue). D. Gene tracks of aligned transcripts over genes that were either upregulated (*Myd88*) or downregulated (*Thbs1*) in multiple depletions. E. Gene ontology analysis of all significantly downregulated genes found in at least 2 or 3 depletion conditions. F. ChromHMM analysis of promoters (500bp upstream of transcription start sites, TSS) of significantly up or downregulated genes found in at least 1, 2, or 3 depletions. Bg corresponds to all genes expressed in primary mouse neurons at DIV10.

We next sought to identify functional enrichment in shared gene sets. We found that genes that were differentially expressed in at least two or three gene sets encoded proteins relevant to neuronal function such as transmembrane transporters and trans-synaptic signaling components (**Fig. 2E**). Interestingly, these genes also shared several chromatin signatures (**Fig. 2F**). Gene sets detected in at least two or three conditions were enriched for genomic regions that contained bivalent domains similar to prior findings (Thudium et al. 2022). These findings suggest that instances of specific genomic features identified in ASD-risk genes (Zhao et al. 2018) (King et al. 2013) are also common features of transcriptional disruptions underlying ASD. Together, these findings indicate that ASD-linked transcriptional regulators share gene targets with chromatin features that may sensitize them to disruption and have common functional outcomes in regulating expression of genes that function at neuronal synapses.

### Gene expression modules influenced by ASD-linked transcriptional regulators

Next, we sought to expand these comparisons using more sensitive measures. We therefore used weighted gene co-expression network analysis (WGCNA) (Zhang and Horvath 2005) to capture system-level changes associated with depletion of ASD-linked transcriptional regulators. We found multiple gene expression modules that were significantly associated with individual ASD-linked transcriptional regulators, as well as modules shared amongst multiple regulators (**Fig. 3A**, **S3A, supplemental data table 4**). In particular, we detected multiple significant modules enriched for genes involved in synapse function and neuronal development, (**Fig 3B-C, S3B**) similar to other analyses (**Fig 1D,2E**). Interesting, we also detected gene modules linked to metabolism (**Fig 3D, S3C**), another major cellular function previously linked to ASD (Rossignol and Frye 2012a, 2012b), and found among upregulated gene sets in other analyses (**Fig S2E**). This suggests that this analysis may reveal more subtle gene expression changes that also contribute to cellular changes associated with ASD. These findings support the broader conclusion that ASD-linked genes have similar functional outcomes rooted in transcriptional signatures that are shared between conditions. Further, these different analyses reveal common trends observed in ASD research more broadly including disruption of neuronal synapses, neuronal development and metabolism.

**Figure 3:**
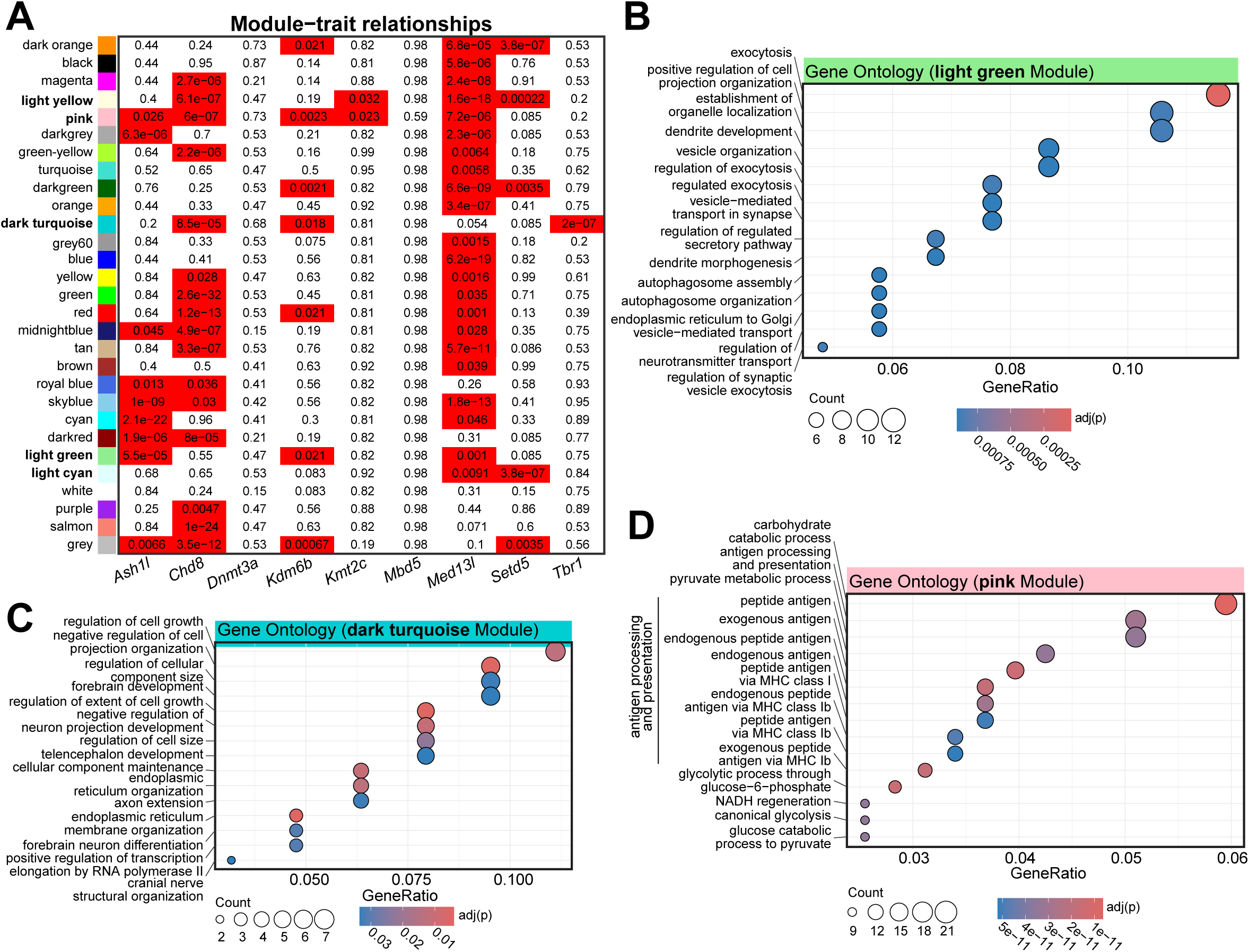
Shared and unique gene modules between ASD linked genes. A. WGCNA modules identified after the co-expression network construction. Numbers are the Benjamini-Hochberg corrected p values for the effect of each depletion on module expression with a linear model. Bolded fonts indicate those with GO terms shown in B-D or in Supplemental Figure 3. B-D. Gene ontology analysis of shared modules. Counts indicate the number of genes in the GO term.

### Defining common transcriptional signatures ASD-linked transcriptional regulators

We next used limma-voom analysis (Law et al. 2014a; Ritchie et al. 2015) to generate a differential gene expression model factoring all transcriptional regulators together to define common transcriptional signatures that diverge from control infected neurons. We found that this multi-condition modeling of the depletions together resulted in significant up and down regulated gene sets with a greater number of downregulated genes identified (**Fig. 4A, Supplemental Data Table 5**). To understand what combinations of transcriptional regulators contributed to the transcriptional signature identified by limma, we examined top hits and found genes with changes in expression across multiple conditions (**Fig. 4B**). Pairwise comparisons of the trends in gene expression of the shared limma signature model showed variable but strong correlations and overlaps between nearly every pair of targets indicating that this approach was successful in identifying convergent gene sets that link all targets (**Fig. 4C**).

**Figure 4:**
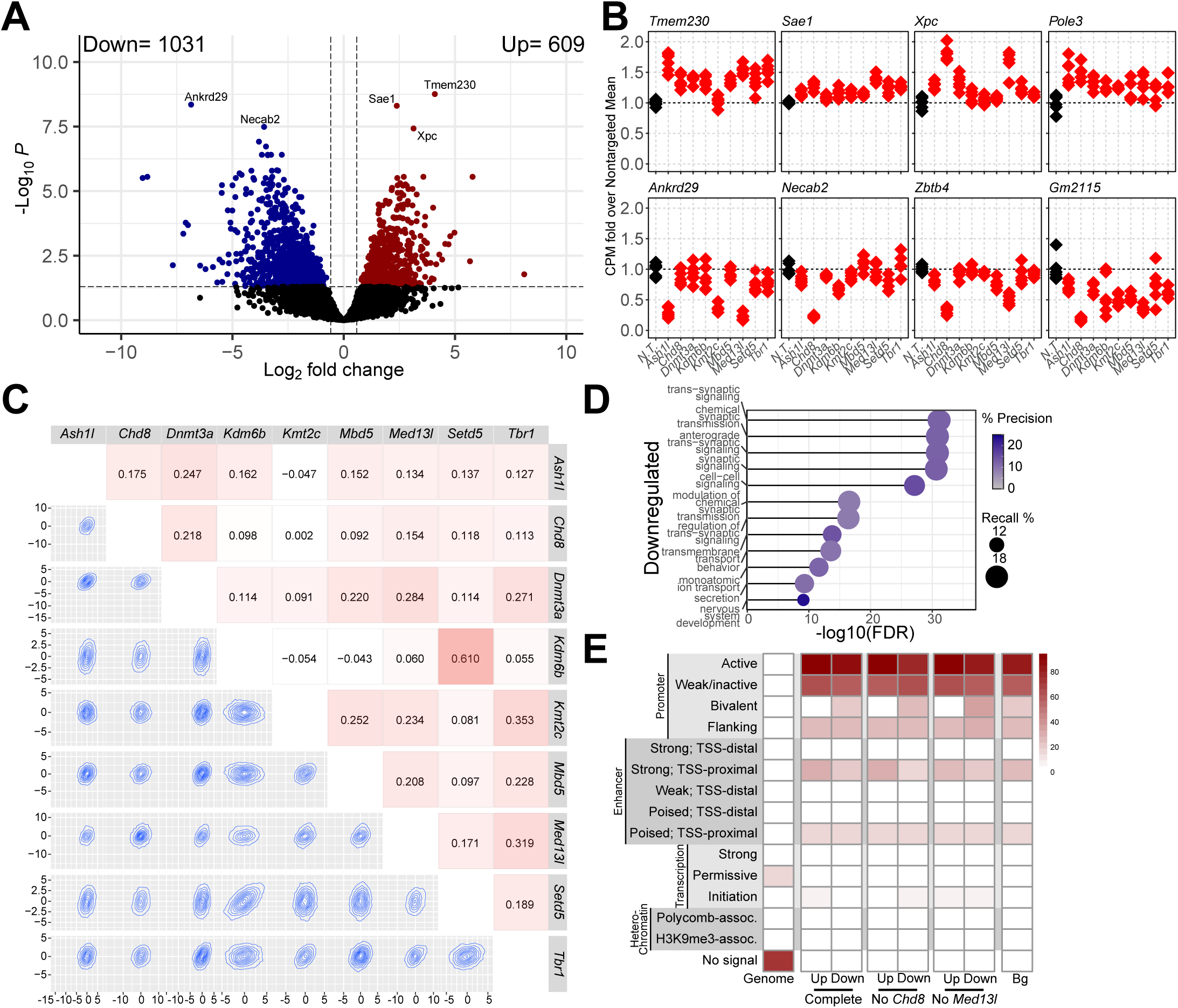
Multiple factor linear modeling with nine independent chromatin modifier depletions in primary mouse neurons. A. Volcano plot of DEGs from the limma/voom model factoring all nine depletion conditions against N.T. control. B. Relative expression of the top DEGs identified in the multi-factor model in each depletion versus the average of nontargeting (N.T.) treated neurons. C. Cross-correlation plots comparing the overall expression profiles of each depletion against N.T. control. Heatmap number and colors indicate correlation coefficient. Topography plots show the distribution of expression changes for all genes between any two depletions within their relative contribution to the model. D. Gene ontology analysis of significantly downregulated DEGs. Recall is the proportion of functionally annotated genes in the query over the number of genes in the GO term. Precision is the number of genes found in the GO term over the total number of genes in the query. E. ChromHMM analysis of promoters (500bp upstream of transcription start sites, TSS) of significantly up or downregulated genes in the complete model, or the models without *Chd8* or *Med13l*. Bg corresponds to all genes expressed in primary mouse neurons at DIV10.

We used gene ontology to examine the function of these gene sets and found that downregulated genes were primarily trans-synaptic signaling while upregulated genes were enriched for those with enzyme activity (**Fig. 4D, S4A**), similar to findings from direct overlap analyses. We again found that downregulated genes are enriched for bivalent domains and upregulated genes are enriched for chromatin features associated with strong proximal enhancers (**Fig. 4E**). We also observed a small but significant overlap between the downregulated signature and ASD and NDD risk genes depending on the dataset used (**Fig. S4B**).

To ensure that gene signatures were not being predominantly driven by single depletion conditions with disproportionately greater effects on gene expression, we repeated these analyses to generate a differential gene expression model without either *Chd8* or *Med13l* depletion conditions, the two targets with the greatest number of DEGs (**Fig. 1C**). We found that in both cases, we detected shared gene expression signatures that encoded similar functional groups and shared most genes with the full model (**Fig. S4C-G**).

Finally, given the recurrent disruption of genes encoding synaptic proteins, regardless of the model or target analyzed, we asked whether synaptic gene expression was sufficient to separate control conditions from depletion conditions. Using a synaptic gene list from GO as an Eigengene vector, we found that this gene set was sufficient to significantly distinguish between control and depletion conditions (**Fig. S5A**). Further, we confirmed that the gene signatures we detected via limma-voom analysis, either using all conditions, or those detected in the absence of *Chd8* or *Med13l*, were similarly able to separate control from depletion conditions as expected (**Fig. S5B-E**).

Together, these findings define a transcriptional signature that is shared across the target genes of multiple ASD-linked transcriptional regulators. This signature expands upon prior work (Thudium et al. 2022) demonstrating that long synaptic genes are highly sensitive to chromatin disruptions. Importantly, these findings demonstrate such features are true of ASD-linked transcriptional regulators more broadly than only ASD-linked chromatin modifying enzymes.

### ASD-risk transcriptional regulators disrupt neuronal firing patterns

We consistently observe considerable disruptions to synaptic gene expression following depletion of both chromatin and transcriptional regulators linked to ASD including in studies that focused on other ASD-linked proteins (Thudium et al. 2022). We therefore asked whether these disruptions resulted in functional changes in neuronal firing patterns. To test this, we used multielectrode array (MEA) recordings to record spontaneous firing activity of neurons depleted for each transcriptional regulator individually (**Fig. 5A, supplemental data tables 6-9**). We began testing at 12 days in culture when spiking activity low but detectable and continued through 22 days in culture. During this period, we detected no changes in covered electrodes or resistance suggesting neurons remained healthy throughout the assay (**Fig. S6A-B**).

**Figure 5:**
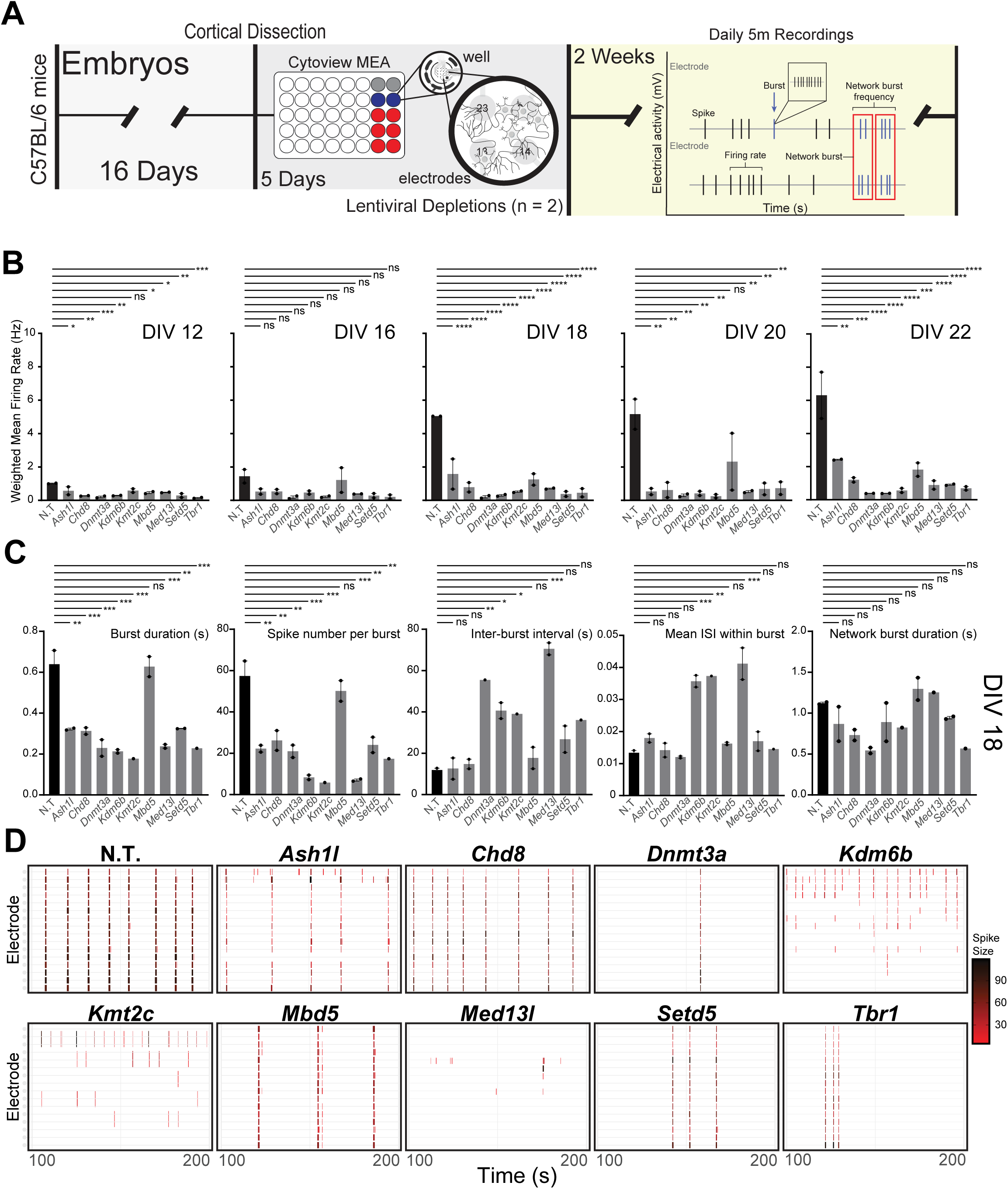
Changes in spontaneous firing activity due to transcriptional regulator depletion. A. Experimental timeline and schematic for the culturing and measurements of primary neurons using the multielectrode array (MEA). The recording diagram illustrates the different metrics collected by the MEA instrument. B. Weighted firing rate of each condition based only on electrodes with activity greater than the minimum spike rate from day in vitro (DIV) 12 to 22. C. Evoked metrics for each condition at DIV 18. Significance based on one-way ANOVA followed by dunnet post-test. *** ≤ 0.001, ** ≤ 0.01, * ≤ 0.05, NS = not significant. D. Example spiking patterns over a 200 second recording period at DIV18.

We found that disruption of every ASD-linked transcriptional regulator tested caused robust changes in firing patterns with decreased spike number over multiple developmental time points (**Fig. 5B, S6C**). Notably, while control neurons show anticipated increases in spiking with maturity, depletion of nearly every transcriptional regulator dampened or fully blocked this increase with maturation. We then focused on 18 days in culture, as a midpoint between early detection of spiking and endpoints when cells begin to show signs of decay after day 22. At day 18, we also observe consistent, strong firing and bursting metrics from our controls allowing us to capture a dynamic range via multiple metrics. Following depletion of multiple transcriptional regulators, we observed decreased bursting and burst duration (**Fig. 5C-D**). Even where bursting was observed, regularity of bursting and synchrony were also severely disrupted (**Fig. 5C, S6D**). Similar though less robust changes were detected at 16 days in culture (**Fig. S6E**).

While depletion of these transcriptional regulators caused many similar functional disruptions to neuronal firing, they also resulted in some divergent effects, particularly for spike amplitude (**Fig. S7A**). While half of the ASD-linked targets showed a decrease in spike amplitude, others showed no change. Further, while *Med13l* depletion caused among the most robust decrease in inter-burst interval and spikes per burst, it also was the only target to show an *increase* in spike amplitude. *Mbd5* loss conversely, caused the least robust effects on multiple metrics (**Fig. 5C**). Given these detectable differences, we sought to group these transcriptional regulators by principal component (PC) analysis based on their effects on neuronal firing. We therefore used all MEA metrics collected and examined PCs one through six and found that PC1 separated for a portion of the ASD-linked targets tested (**Fig. S7B**). We therefore repeated lima voom analysis on the groups that most clearly separated based on MEA metrics, comparing *Tbr1, Dnmt3a* and *Med13l* to *Chd8, Ash1l,* and *Mbd5* (**Fig. S7C**). We found that comparisons between these subgroups identified significant gene expression changes with enriched functions relating to metabolism (**Fig. S7D**). This suggests that divergent transcriptional disruptions may underly functional differences between various ASD-linked genes in addition to their shared disruptions causing numerous overlapping effects. Together, these data provide evidence that a variety of gene disruptions linked to ASD are capable of severely disrupting neuronal firing and that downregulation of synaptic gene expression detected via RNA-sequencing results in robust functional changes. Further, these findings provide evidence of convergent functional effects while also highlighting the subtle but detectable differences between ASD-linked transcriptional regulators.

## Discussion

Here, we partially depleted chromatin regulators, DNA modifying enzymes, and transcription factors that are all strongly linked to ASD. We found that despite divergent functions, they converge on a common gene expression signature identified through multiple analyses. Using RNA-sequencing, we defined gene expression signatures common between these ASD-linked transcriptional regulators. This signature encodes critical neuronal proteins including synaptic proteins suggesting similar functional outcomes of these transcriptional disruptions. To test the functional implications of these transcriptomic changes, we measured neuronal firing patterns. Every transcriptional regulator tested affecting spiking and bursting patterns throughout neuronal maturation. These data demonstrate that disruption of multiple types of ASD-linked transcriptional regulators results in shared gene expression signatures and neuronal firing patterns despite having divergent effects on chromatin and gene regulation.

This work expanded upon prior findings by testing transcriptional regulators broadly rather than just focusing on single ASD-linked genes or on only proteins that directly modify chromatin. Instead, we selected proteins with highly divergent effects on chromatin and transcription. These included chromatin modifying enzymes that target different histone sites, chromatin remodelers from different complexes, a DNA modifying enzyme, a transcription factor, and a non-catalytic chromatin-complex protein. Even with this expanded group, we successfully identified both shared gene signature and common neuronal spiking and bursting patterns. This suggests that specific neuronal gene sets are particularly sensitive to multiple types of epigenetic manipulations and that disruption of these genes causes robust functional changes in neuronal firing.

Both the analyses overlapping individual depletion DEGs and the limma model identified several significantly downregulated genes that may be important points of convergence in the shared roles of the modifiers in affecting neuronal functioning. Topoisomerase IIα (Top2a) was downregulated in 7 of the 9 depletions in addition to being identified in the limma model. Notably, loss of Top2a causes social deficits in mice and zebrafish (Geng et al. 2022) suggesting its decreased expression may have functional consequences that are relevant to ASD phenotypes. Additionally, both analyses identified numerous genes important in brain development including Insulin-like growth factor 1 and 2 (Igf1/2) and Neuronal PAS Domain Protein 1 (Npas1). We also observed consistent reduction in two aquaporins (Aqp4 and Aqp11) that are less well studied but also linked to social and anxiety-like behavior in animal models (Davoudi et al. 2023). Together with the overall reductions found in the expression data by GSEA and the functional results of the MEA, these findings support the convergent roles of these chromatin modifiers on neuronal biology that may contribute to neurodevelopmental disorders. WGCNA analysis similarly identified several relevant modules including those enriched for genes involved in cellular respiration and metabolism. Notably, metabolic dysfunction is also linked with ASD (Rossignol and Frye 2012a, 2012b) and may contribute to neuronal aberrations in the disorder due to the high energy demands of neurons.

While we identified numerous genes of interest, the system used here also has several limitations. The signatures we defined are likely to be somewhat specific to the developmental time selected and the model chosen. We expect that if these assays were performed at different developmental time points, we would detect different gene expression signatures that may have very distinct effects on functional outcomes. We thus view these results as primarily indicative of sensitive genes during this stage of neuronal maturation rather than an absolute list of transcriptional targets for any given gene studied here. In addition, it is likely that results will at least partly diverge from human systems such as iPSC derived neurons that may more closely match human developmental trajectories. Nonetheless, we view this system as offering unique insights that emerge from the ability to test multiple ASD risk genes in parallel in a highly controlled system with genetically identical biological replicates.

Together our findings demonstrate shared gene expression signatures and functional outcomes of disruption of multiple ASD-linked transcriptional regulators. This work provides insights into underlying cellular mechanisms that link ASD-risk genes to neuronal function and highlight common pathways that are controlled by ASD-linked transcriptional regulators.

## Methods

### Mice

All mice used were on the C57BL/6J background, housed in a 12 hour light–dark cycle, and fed a standard diet. All experiments were conducted in accordance with and approval of the IUCAC.

### Primary Neuronal Culture

Cortices were dissected from E16.5 C57BL/6J embryos and cultured in supplemented neurobasal medium (Neurobasal [Gibco 21103-049], B27 [Gibco 17504044], GlutaMAX [Gibco 35050-061], Pen-Strep [Gibco 15140-122]) in TC-treated 12 or 24 well plates coated with 0.05 mg/mL Poly-D-lysine (Sigma-Aldrich A-003-E). Neurons from individual embryos were seeded to 12 wells each. At 3-4 DIV, neurons were treated with 0.5 mM AraC. For all experiments using cultured cortical neurons, neurons were treated with lentivirus containing shRNA on DIV 5. For RNA and Protein analysis cells were collected at DIV 10.

### Virus Generation and validation

HEK293T cells were cultured in high-glucose DMEM growth medium (Corning 10-013-CV), 10% FBS (Sigma-Aldrich F2442-500ML), and 1% Pen-Strep (Gibco 15140-122). Calcium phosphate transfection was performed with Pax2 and VSVG packaging plasmids in serum-free media. shRNAs in a pLKO.1-puro backbone were purchased from the Sigma-Aldrich Mission shRNA library (SHCLNG) and shown in **Supplemental Table 11**. Media was changed 2h after transfection and viruses were collected 24 and 48hr later. Viral media was passed through a 0.45-μm filter and precipitated overnight with PEG-it solution (40% PEG-8000 [Sigma-Aldrich P2139-1KG], 1.2 M NaCl [Fisher Chemical S271-1]). Viral particles were pelleted at 1500g, washed, and resuspended in 200 μL PBS. Virus Validation was done on E16.5 WT cortical neurons. Neurons were infected with virus on DIV 5. Depletion efficiency was measured using RT-qPCR with shRNA targeting luciferase as a non-targeting control.

### Western

After DIV 10, neurons were lysed in RIPA (25 mM Tris at pH 7.6, 150 mM NaCl, 1% NP-40, 1% sodium deoxycholate, 0.1% SDS) supplemented by protease inhibitor (Roche 04693124001), phosphatase inhibitor (Roche 04906837001), 1mM DTT, 1mM PMSF. Lysates were mixed with 5X Loading Buffer (5% SDS, 0.3 M Tris pH 6.8, 1.1 mM Bromophenol blue, 37.5% glycerol), boiled for 10 minutes, sonicated for 10 minutes, and cooled on ice. Protein was resolved by 4– 20% Tris-glycine or 3–8% Tris-acetate SDS-PAGE (invitrogen novex gels XP04205, EA03785) followed by transfer to a 0.45-μm PVDF membrane (Sigma-Aldrich IPVH00010) for immunoblotting. Membranes were blocked for 1 hour in 1-5% BSA or 5% milk in TBST and probed with primary antibody overnight at 4C. Antibodies are shown in **Supplemental Table 12**.

### RNA Isolation

Total RNA was collected from all cultures at DIV 10 using the Zymo Quick-RNA microprep kit (R1050).

### RT-qPCR

250ng of RNA per sample was used to prepare cDNA using the high-capacity cDNA reverse transcription kit (Applied Biosystems 4368813), and quantitative PCR was performed with Power SYBR Green PCR master mix (Applied Biosystems 4367659). Primers are shown in **Supplemental Table 13**.

### RNA-seq Library Preparation

Sequencing libraries were prepared using the TruSeq Stranded mRNA kit (Illumina 20040534). Prior to sequencing, library size distribution was confirmed by capillary electrophoresis using an Agilent 4200 TapeStation with high sensitivity D1000 reagents (5067-5585), and libraries were quantified by qPCR using a KAPA Library Quantification Kit (Roche 07960140001). Libraries were sequenced on an Illumina NextSeq1000 instrument (100-bp read length, paired end).

### Data Processing and Differential Gene Expression

Reads were mapped to *Mus musculus* genome build mm10 with STAR(Dobin et al. 2013) (v2.7.1a) and assigned to exonic features using the featureCounts function of subRead (v2.0.3). DESeq2(Love et al. 2014) (v1.38.0) was used for pairwise differential gene expression analysis using a negative binomial model with default model fitting parameters. 6 WT replicates were used for downstream analysis for each condition. Batch effects introduced by differences in the sexes were regressed using a negative binomial model via ComBat-ref (Zhang 2024). Subsetting filters were applied to determine significant DEGs as follows: baseMean of normalized counts > 20, adjusted p value ≤ 0.05, and |fold change| ≥ 1.5 (|log_2_(FC)| ≥ 0.58). Integrated Genomics Viewer (Robinson et al. 2011) (v2.16.0.01) was used to visualize RNA-seq read track that were normalized to the same total read number across all samples using SAMtools. Multifactor gene expression analysis was performed using limma (v3.52.4) via edgeR(Robinson et al. 2010) (v3.38.4). Normalization factors were calculated, and expression values were transformed to logCPM using voom(Law et al. 2014b). An empirical Bayesian distribution was calculated with weighting from all 9 depletion conditions across all samples.

### Downstream Analysis

Hypergeometric testing of significant overlaps between gene lists was performed using the dhyper density distribution without replacement using a background size of 13713 and Bonferroni correction for multiple testing. gProfiler2(Kolberg et al. 2020) (v0.2.3) was used for gene ontology analysis with g:SCS correction of multiple testing, a background list of all genes expressed in the neuronal culture (base mean of normalized counts > 20), and a term size limit of 2000 genes. SynGO was performed using the web client(Koopmans et al. 2019) (v1.2). Gene set enrichment analysis was performed using FGSEA (Korotkevich et al. 2016) (v1.22.0). This utilized a multilevel Monte Carlo approach with default parameters. Gene lists were from REACTOME or SynGO synapse umbrella terms. ChromHMM (Ernst and Kellis 2017) analysis used validated ChIP-seq data from E12.5 mouse forebrain tissue against 500bp windows directly upstream of query genes. Transcription factor motif analysis on 5kb windows directly upstream of query genes were analyzed using sequence enrichment analysis in the MEME suite(Bailey and Grant 2021) (v5.5.7). Motifs were scanned using the JASPAR 2020 Core Vertebrates database using the windows 5kb upstream of 1000 randomly selected genes expressed in neurons. WGCNA was used to construct a signed co-expression network(Langfelder and Horvath 2008) (v1.73). Batch-corrected inputs were filtered for those with read count ≥ 10 and normalized using variance stabilizing transformation of DESeq2 with design = ∼ depletion. Network construction used a soft threshold blcokSize = 20000, networkType = “Signed”, power of 20, scale free topology fit of 0.83, and minimum module size of 40. For each module, we calculated its expression (ME, module eigengene) as the first principal component of normalized expression. We used a linear model to test the effect of each depletion on module expression compared to non-targeting control. P values were corrected using Benjamini-Hochberg correction. Gene ontology for WGCNA was performed using the enrichGO function from clusterProfiler (Yu et al. 2012) (v4.10.1) with the BP ontology database, Benjamini-Hochberg correction, p value cutoff of 0.05, and q value cutoff of 0.05. The top 15 terms per analysis were selected for figures.

### MEA Analysis

Primary neuronal culturing was performed as described above using E16.5 embryos derived from mating of Female mice (C57BL/6N). Cerebral cortices were isolated from two pups from two separate litters and dissociated as described above, then plated in poly-D-lysine/laminin-coated multielectrode array plates (M768-tMEA-48W, Axion Biosystems) using the dot plate method, at 70K cells/well. For viral infections, each well was uniquely infected with one of the 9 ASD targets or non-targeting shRNA at DIV 5. Viral media was removed after 20 hr of exposure and replaced with enriched neurobasal media. After 11 days, the synaptic activity of cultured neurons was recorded daily using the Maestro MEA system (Axion Biosystems, Atlanta, GA, USA). Recordings were done 5 and 30 minutes after placing in MEA machine at 5% CO_2_ and at 37C. Metrics were further processed using Prism with a one-way ANOVA followed by dunnet post-test and R.

## Data access

All raw and processed sequencing data generated in this study have been submitted to the NCBI Gene Expression Omnibus (GEO; https://www.ncbi.nlm.nih.gov/geo/) under accession number **GSE286067**.

## Competing Interests

The authors have no conflicts of interest.

## Acknowledgments

E.K and R.A. were supported by NIH NIMH (1DP2MH129985) and NIH NINDS (R01NS134755), Autism Spectrum Program of Excellence at the University of Pennsylvania (ASPE), and the Eagles Autism Foundation. A.P. was supported by NIH grant T32-ES019851. JEPC was supported by NIH NIMH (1R01MH120269; 1DP1MH129957); NIH NINDS (5-R01-NS114226). S.S. was supported by University of Pennsylvania CURF grants. S.Z. was supported by the University of Pennsylvania Epigenetics Institute. We acknowledge the use of micro-electrode array equipment purchased through use of a NIH Shared Equipment Grant (1S10OD032363 to J.E.P-C and E.K.). R.A. performed cell culture and MEA assays and generated RNA-sequencing libraries. A.P. Analyzed sequencing and MEA data. S.S. provided analysis support for MEA data. S.Z provided WGCNA analysis. J.E.P-C and A.J.W. provided MEA access and guidance. E.K. wrote the manuscript and led the project.

## Notes

### Competing Interest Statement

The authors have declared no competing interest.

### Summary of Updates

Author information, new experiments, new analysis.

