## Supplemental Material for "Autism spectrum disorder risk genes have convergent effects on transcription and neuronal firing patterns in primary neurons"

**Supplemental Figure 1: Validation of depletions.**

**Supplemental Figure 2: Transcriptomic and overlapping features of individual chromatin modifier loss.**

**Supplemental Figure 3: WGCNA analysis module gene ontology.**

**Supplemental Figure 4: Multiple factor linear modeling omitting *Chd8* or *Med13l*.**

**Supplemental Figure 5: Eigengenes of broad transcriptional signatures separate controls from modifier depleted neurons.**

**Supplemental Figure 6: Multi-electrode array quality control and additional metrics.**

**Supplemental Figure 7: Multi-electrode array differences between depletions and control reveals subgroups.**

**Supplemental Data Table 1. Differential gene expression by depletion.**

**Supplemental Data Table 2. Gene ontology results by depletion.**

**Supplemental Data Table 3. Overlap of DEGs.**

**Supplemental Data Table 4. WGCNA modules.**

**Supplemental Data Table 5. Limma voom analysis of depletions.**

**Supplemental Data Table 6. MEA mean firing rate.**

**Supplemental Data Table 7. MEA resistances.**

**Supplemental Data Table 8. MEA weighted mean firing rates.**

**Supplemental Data Table 9. MEA DIV16 and DIV18 metrics.**

**Supplemental Data Table 10. MEA metric groups RNA-seq comparisons.**

**Supplemental Data Table 11. shRNA oligo sequences.**

**Supplemental Data Table 12. Antibodies and dilutions used.**

**Supplemental Data Table 13. Primer sequences.**

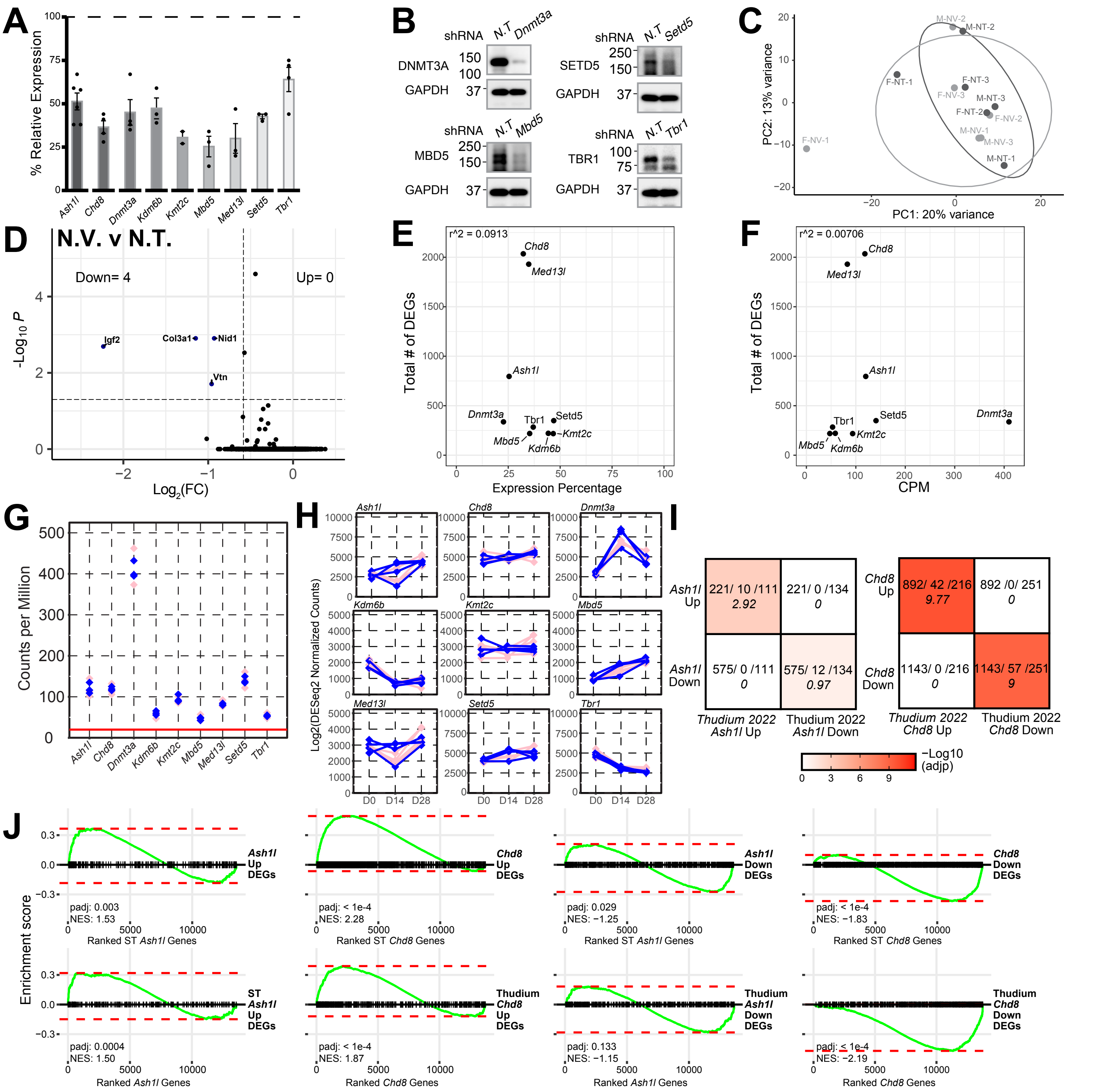

**Supplemental Figure 1: Validation of depletions.** A. Relative transcript expression level of each chromatin modifier following depletion versus non-targeting control virus (N.T.) treated neurons as measured by RT-qPCR. B. Western blots showing protein level of four of the chromatin modifiers in control-treated and shRNA-treated neurons. C. Principal component analysis of the pairwise comparison between neurons with no virus (N.V.) and those treated with N.T. control (n = 6, 3 female and 3 male for each condition). D. Volcano plot of differential gene expression between no N.V. and N.T. neurons. E-F. Scatterplot of the number of DEGs identified in the comparison between a depletion of a modifier and control against the average expression of that modifier following depletion (E) or the counts per million (CPM) normalized expression of that modifier (F). G. Counts per million of each chromatin modifier in DIV10 primary mouse neuron culture. Each replicate is colored by sex (female: pink, male: blue). The red line indicates the expression noise threshold of 20. H. DESeq2 normalized expression counts of each chromatin modifier in cortical excitatory neurons (Nex+) isolated from mice at 0, 14, and 28 days of age from GEO:GSE282467. I. Overlap of significantly up or downregulated genes in the depletion of Ash1l or Chd8 in this dataset (rows) with the same depletions in Thudium 2022. (DEGs found in this dataset / overlap / DEGs found in Thudium 2022) and hypergeometric test results are shown for each overlap. J. GSEA of the change in expression of up and downregulated genes in this dataset across the Thudium 2022 dataset (top) or DEGs in Thudium 2022 across this dataset (bottom), for either the depletion of Ash1l or Chd8. Each dark line corresponds to the location of a DEG is in the expression rank ordered list of the other dataset.

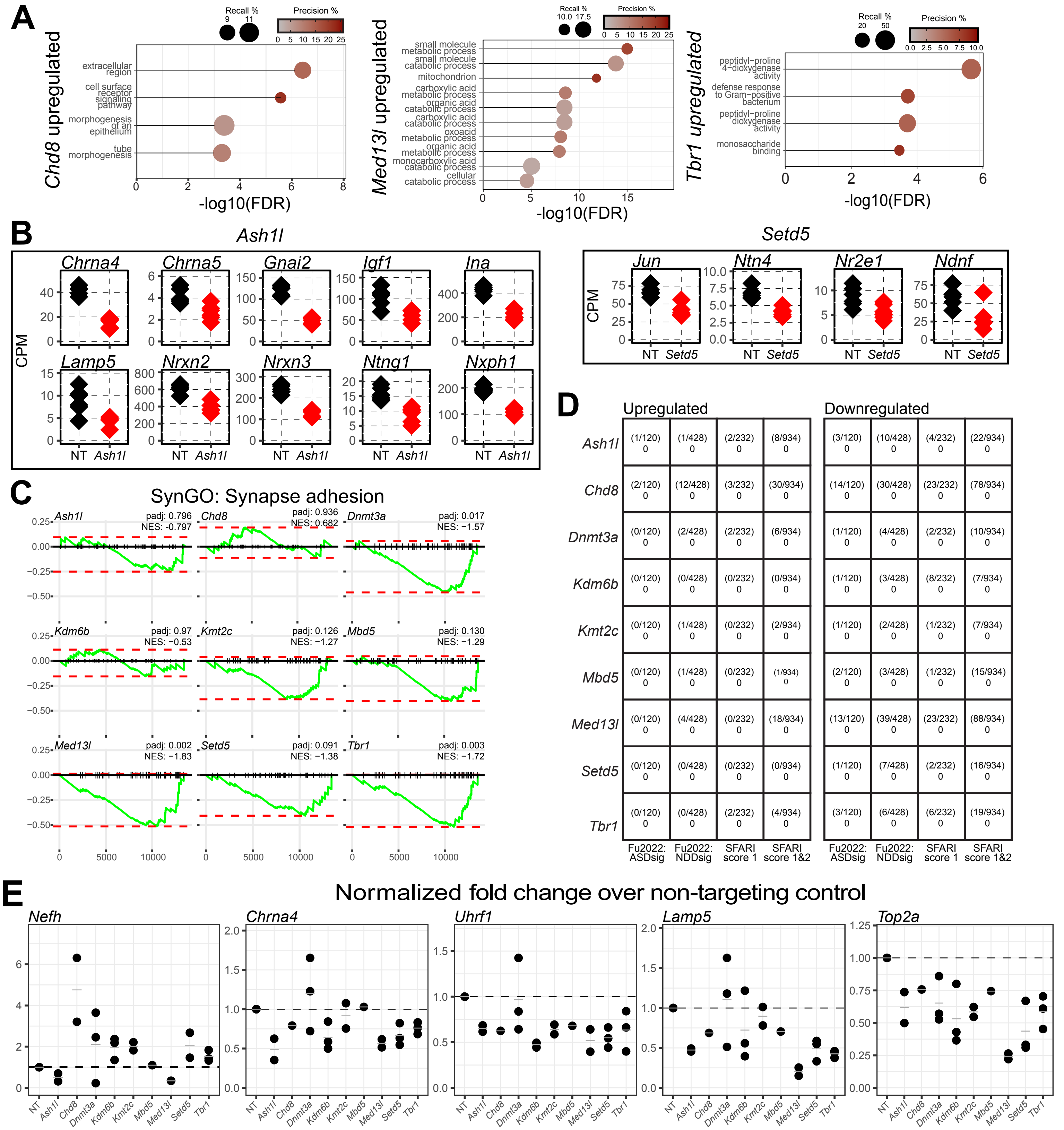


**Supplemental Figure 2: Transcriptomic and overlapping features of individual chromatin modifier loss.** A. Gene ontology analysis of differentially upregulated genes for the depletions of Med13l, Tbr1, or Chd8. B. Counts per million normalized expression of differentially downregulated genes encoding proteins involved in postsynaptic signaling for N.T. control or each depletion of Ash1l and Setd5. C. GSEA of genes in the SynGO synapse adhesion term for each depletion transcriptional signature. The green line is the running, normalized enrichment score across all genes ranked by their change in expression from most increased at 1 to least increased. The red dotted lines specify the maximum and minimum running enrichment score per depletion. NES indicates the directional normalized enrichment score across all genes. D. Overlap of significantly up or downregulated genes for each depletion with ASD or NDD associated genes from Fu et al (FDR < 0.05), or the SFARI database for autism-associated genes with a score of 1 or 1&2 (DEGs found in the external list / total number of genes in the external list) and hypergeometric test results are shown for each overlap. E. Gapdh normalized qPCR fold differences for candidate genes in the depletions of each modifier against non-targeting (N.T.) control for increased (red) and decreased (blue) genes.

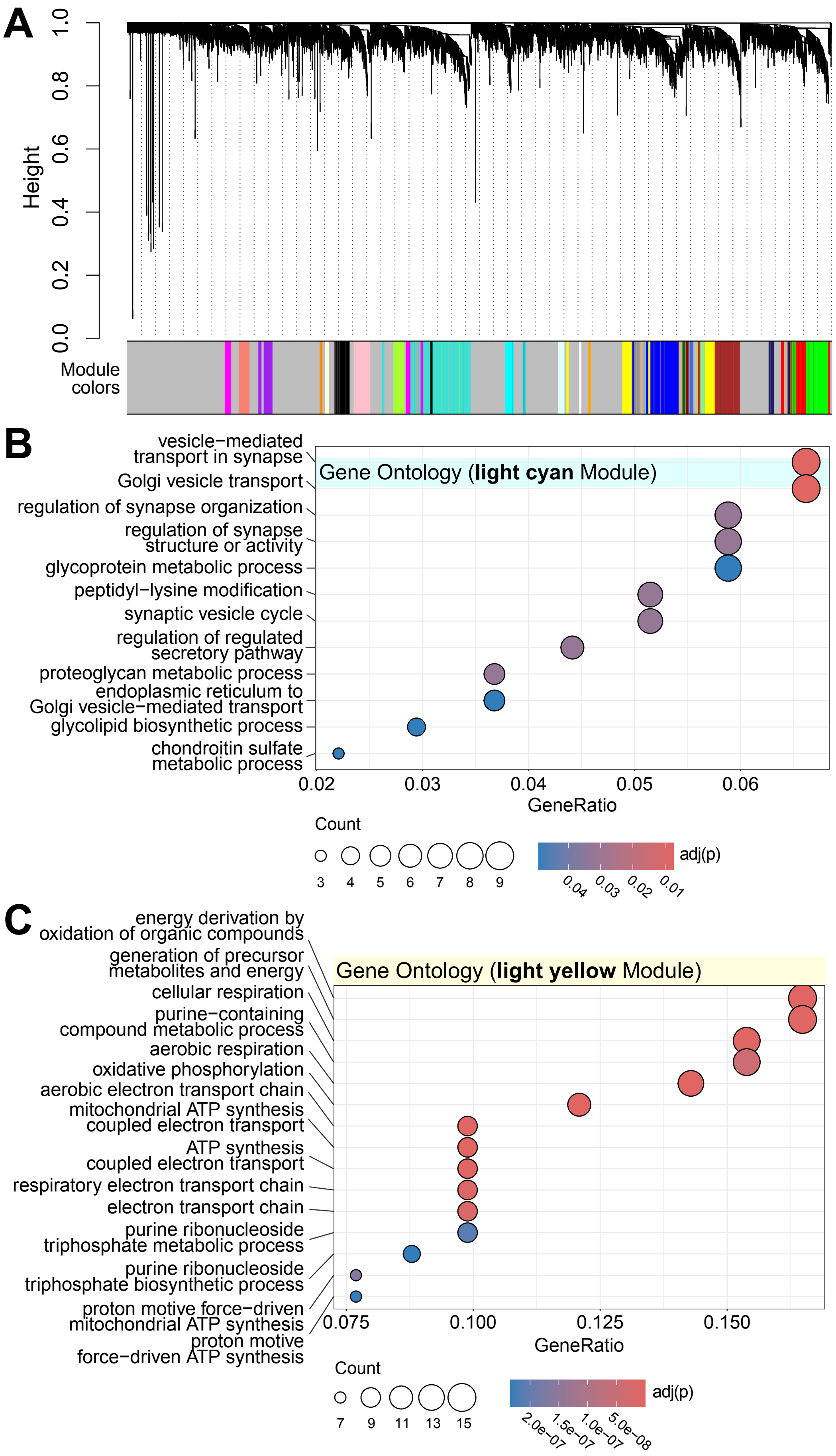


**Supplemental Figure 3: WGCNA analysis module gene ontology.** A. Cluster dendrogram of the classification of genes to each model. B-C. Gene ontology analysis of shared modules.

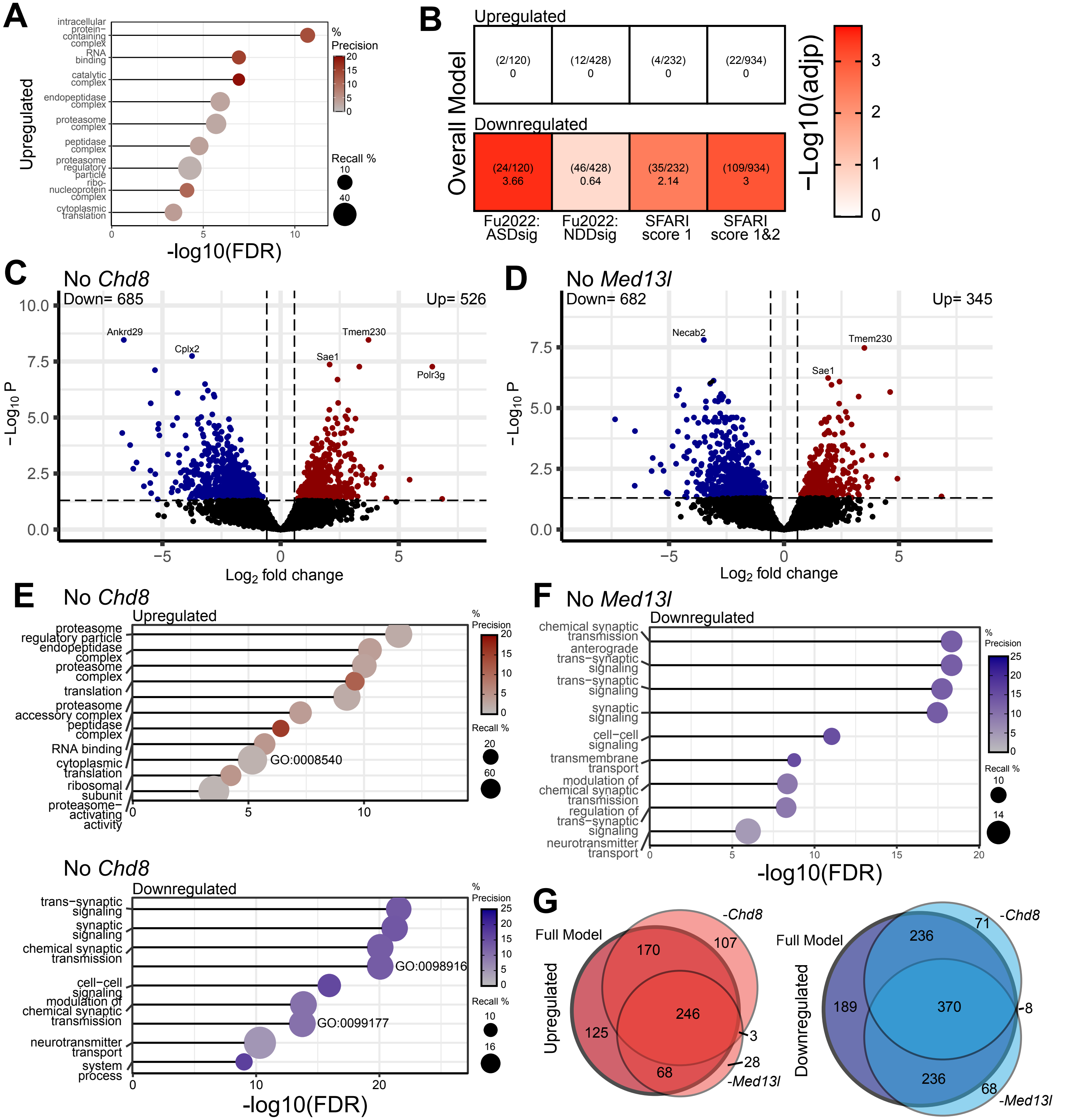


**Supplemental Figure 4: Multiple factor linear modeling omitting Chd8 or Med13l.** A. Gene ontology analysis of significantly upregulated DEGs. Recall is the proportion of functionally annotated genes in the query over the number of genes in the GO term. Precision is the number of genes found in the GO term over the total number of genes in the query. B. Overlap of significantly up or downregulated genes from the complete limma/voom model with ASD or NDD associated genes from Fu et al (FDR < 0.05), or the SFARI database for autism-associated genes with a score of 1 or 1&2. DEGs found in the external list / total number of genes in the external list (in parentheses) and hypergeometric test results are shown for each overlap. C-D. Volcano plots of DEGs from the limma/voom models factoring eight depletion conditions against N.T. control without either Chd8 (C) or Med13l (D). E-F. Gene ontology analysis of significantly upregulated and downregulated DEGs in the limma/voom model without Chd8 (C) and downregulated DEGs in the model without Med13l (F). No significant terms were identified for -Med13l model upregulated genes. G. Venn diagrams of the overlap between differentially up or downregulated genes from the complete model and the models omitting Chd8 or Med13l.

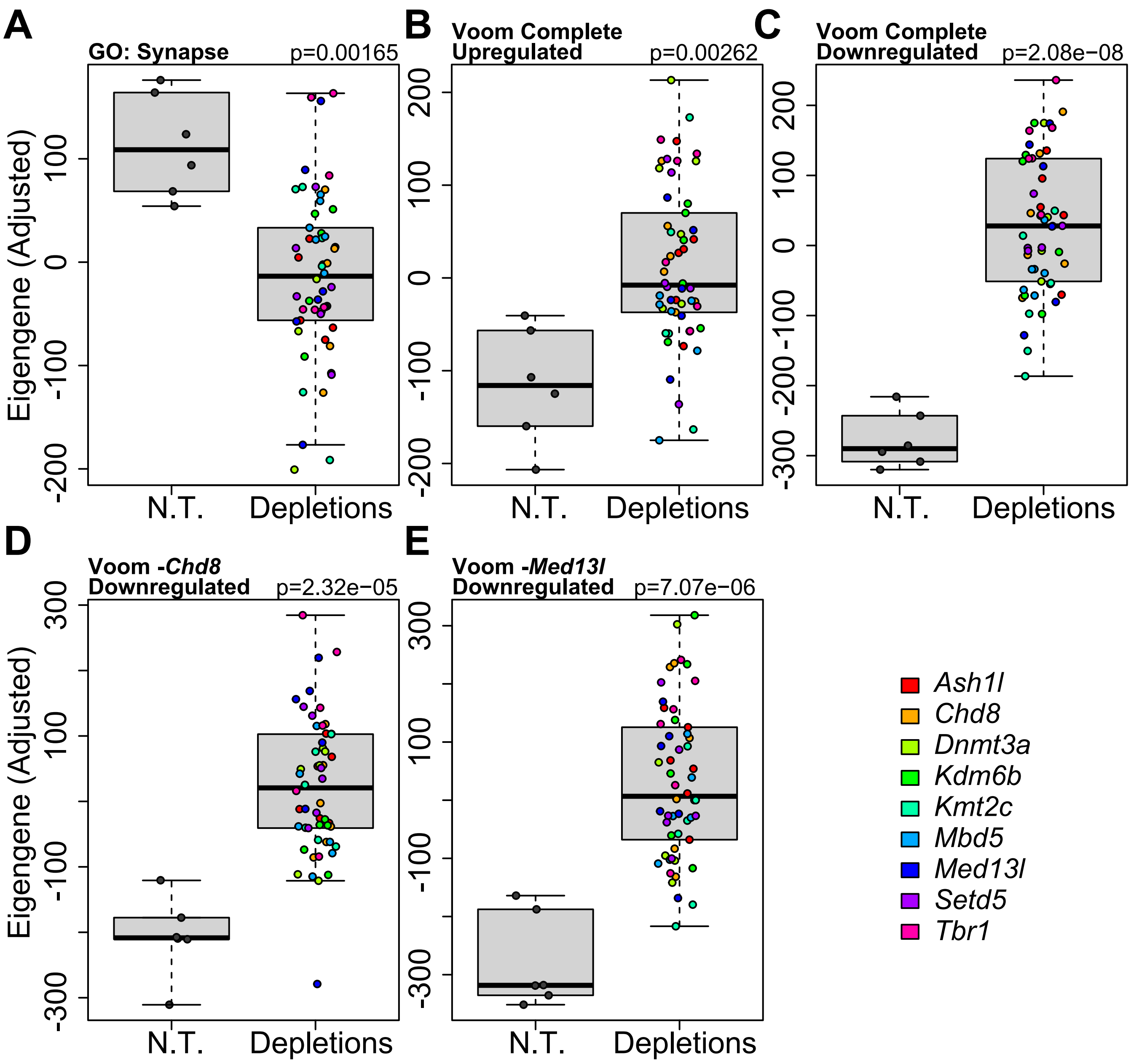


**Supplemental Figure 5: Eigengenes of broad transcriptional signatures separate controls from modifier depleted neurons.** A-E. Eigengene separation of non-targeting (N.T.) controls and aggregated depletion profiles using the genes from the GO synapse term GO:0045202 (A), significantly upregulated genes from the limma/voom complete model (B), significantly downregulated genes from the same model (C), or significantly downregulated genes from the model without *Chd8* (D) or *Med13l* (E). Note that Eigengene values (y axis) do not equate to the direction (up or down) of gene expression changes.

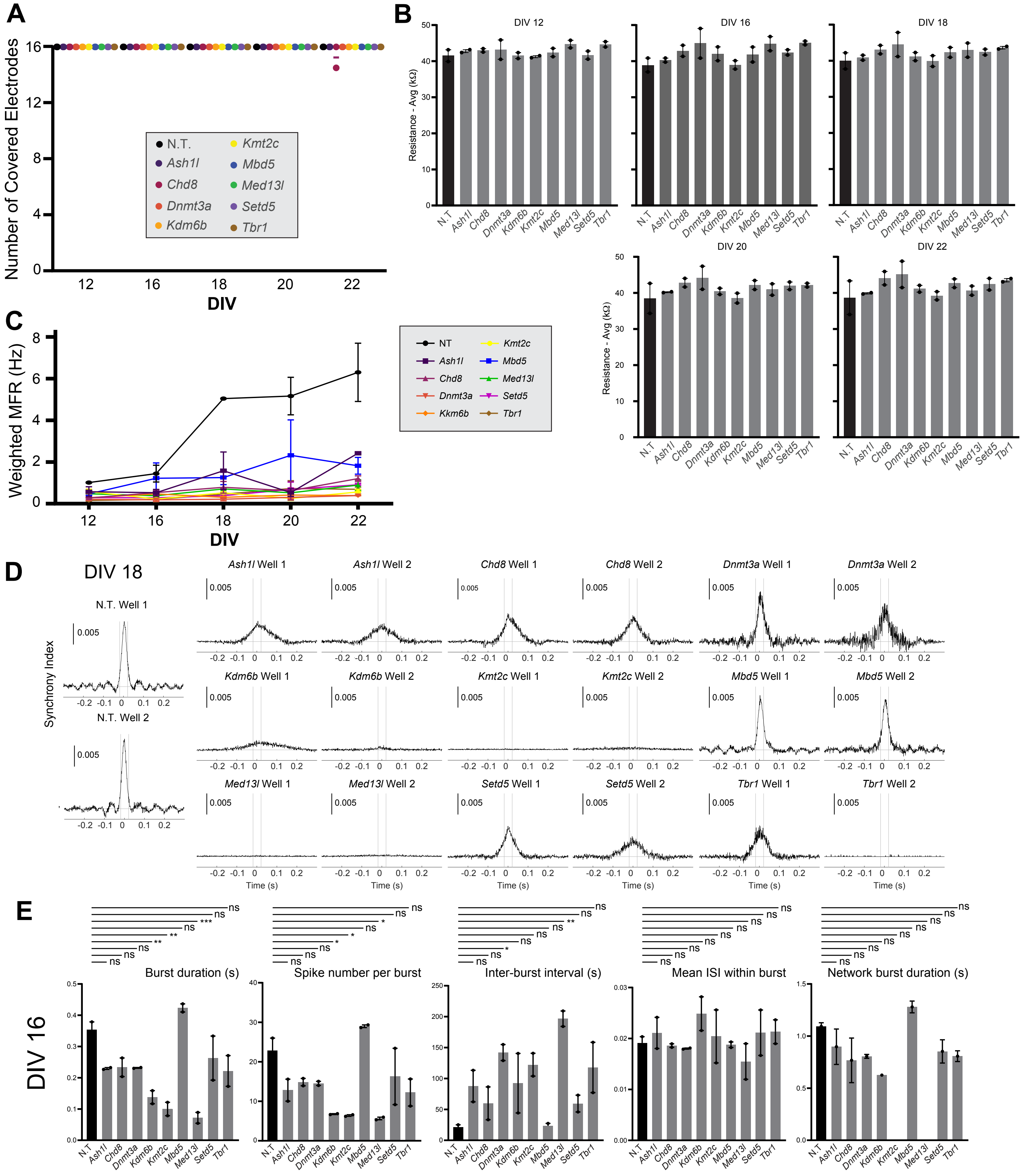


**Supplemental Figure 6: Multi-electrode array quality control and additional metrics.** A. Number of covered electrodes per well per condition as defined by those with resistance greater than the threshold of 18 kΩ. B. Average resistance in each well across conditions from day in vitro (DIV) 12 to 22. All comparisons are non-significant. C. Weighted firing rate of each condition from DIV 12 to 22. D. Synchrony measures for each well of each condition at DIV 18 as defined the inter-electrode cross correlation across all spikes. E. Metrics for each condition at DIV 16. Significance based on one-way ANOVA followed by dunnet post-test. *** ≤ 0.001, ** ≤ 0.01, * ≤ 0.05, NS = not significant. Hz indicates hertz. MFR indicates mean firing rate. ISI indicates interspike interval. S indicates seconds

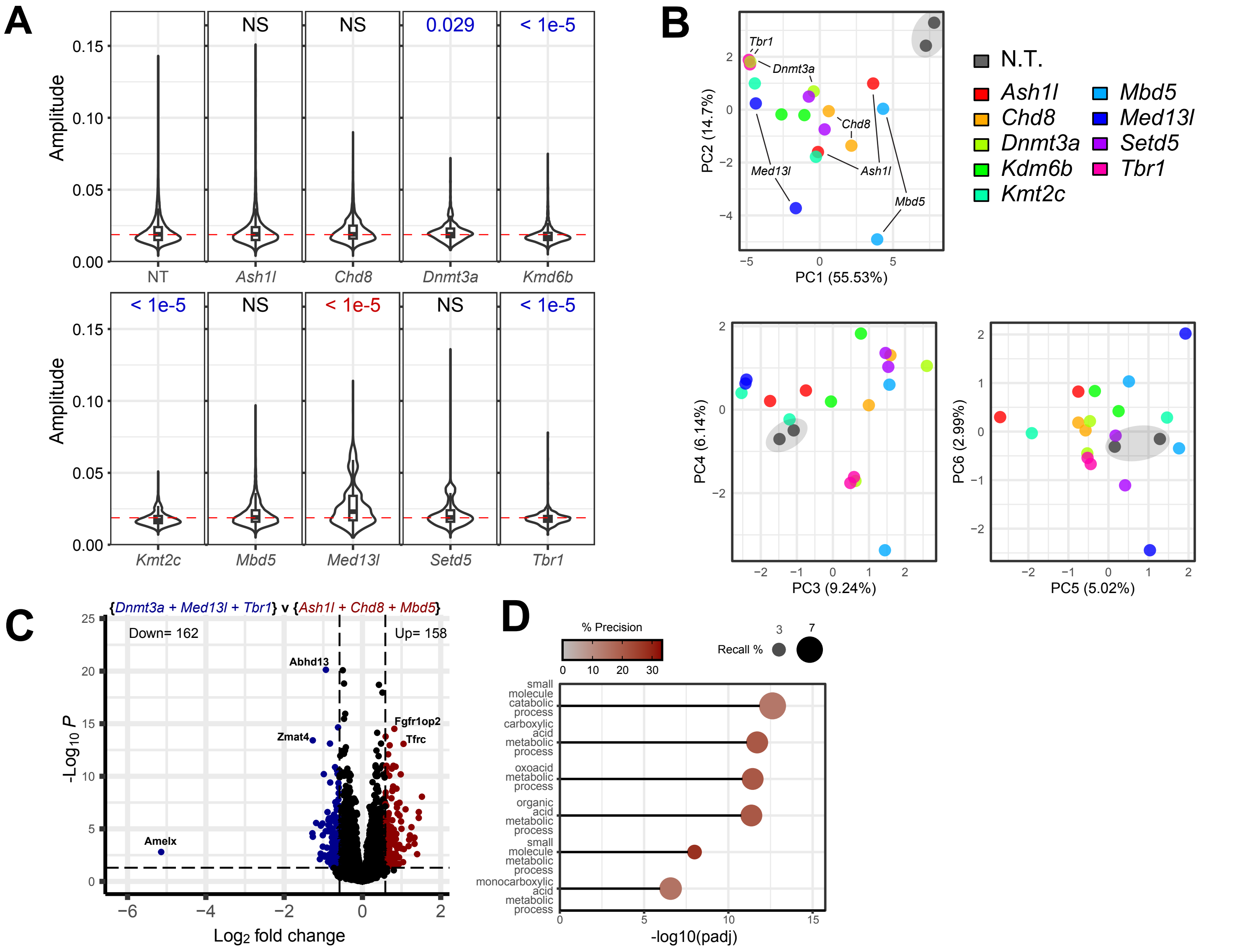


**Supplemental Figure 7: Multi-electrode array differences between depletions and control reveals subgroups.** A. Violin plots of the aggregated spike amplitudes between each two replicates per condition over the entire 300 second run on day in vitro (DIV) 18. The red line indicates the mean of non-targeting (N.T.) control spikes. Significance based on two-sided T-tests against N.T. control are shown above. NS = not significant. Text is colored blue or red if the means are lower or greater than N.T., respectively. B. Principal component analysis of each condition replicate using all MEA average metrics calculated on DIV 18 for the corresponding well. Metrics and values are shown in supplemental table 9. Significance based on one-way ANOVA followed by dunnet post-test. C. Volcano plot of differentially expressed genes in the pairwise comparison of the aggregate profiles of *Dnmt3a/Med13l/Tbr1* against *Ash1l/Chd8/Mbd5*. D. Gene ontology analysis of genes significantly greater in the *Ash1l/Chd8/Mbd5* (or lower in *Dnmt3a/Med13l/Tbr1*) group.
